# Structural models of SARS-CoV-2 Omicron variant in complex with ACE2 receptor or antibodies suggest altered binding interfaces

**DOI:** 10.1101/2021.12.12.472313

**Authors:** Joseph H. Lubin, Christopher Markosian, D. Balamurugan, Renata Pasqualini, Wadih Arap, Stephen K. Burley, Sagar D. Khare

## Abstract

There is enormous ongoing interest in characterizing the binding properties of the SARS-CoV-2 Omicron Variant of Concern (VOC) (B.1.1.529), which continues to spread towards potential dominance worldwide. To aid these studies, based on the wealth of available structural information about several SARS-CoV-2 variants in the Protein Data Bank (PDB) and a modeling pipeline we have previously developed for tracking the ongoing global evolution of SARS-CoV-2 proteins, we provide a set of computed structural models (henceforth models) of the Omicron VOC receptor-binding domain (omRBD) bound to its corresponding receptor Angiotensin-Converting Enzyme (ACE2) and a variety of therapeutic entities, including neutralizing and therapeutic antibodies targeting previously-detected viral strains. We generated bound omRBD models using both experimentally-determined structures in the PDB as well as machine learningbased structure predictions as starting points. Examination of ACE2-bound omRBD models reveals an interdigitated multi-residue interaction network formed by omRBD-specific substituted residues (R493, S496, Y501, R498) and ACE2 residues at the interface, which was not present in the original Wuhan-Hu-1 RBD-ACE2 complex. Emergence of this interaction network suggests optimization of a key region of the binding interface, and positive cooperativity among various sites of residue substitutions in omRBD mediating ACE2 binding. Examination of neutralizing antibody complexes for Barnes Class 1 and Class 2 antibodies modeled with omRBD highlights an overall loss of interfacial interactions (with gain of new interactions in rare cases) mediated by substituted residues. Many of these substitutions have previously been found to independently dampen or even ablate antibody binding, and perhaps mediate antibody-mediated neutralization escape (*e.g*., K417N). We observe little compensation of corresponding interaction loss at interfaces when potential escape substitutions occur in combination. A few selected antibodies (*e.g*., Barnes Class 3 S309), however, feature largely unaltered or modestly affected protein-protein interfaces. While we stress that only qualitative insights can be obtained directly from our models at this time, we anticipate that they can provide starting points for more detailed and quantitative computational characterization, and, if needed, redesign of monoclonal antibodies for targeting the Omicron VOC Spike protein. In the broader context, the computational pipeline we developed provides a framework for rapidly and efficiently generating retrospective and prospective models for other novel variants of SARS-CoV-2 bound to entities of virological and therapeutic interest, in the setting of a global pandemic.

## Introduction

The emerging Omicron Variant of Concern (VOC) (B.1.1.529) of SARS-CoV-2, first sequenced in the Republic of South Africa, has recently been deemed as Variant of Concern (VOC) by the World Health Organization (WHO) and is currently spreading within the human population worldwide.^1^ However, at the time of writing in December 2021, relatively little is known about its transmissibility, potential for immune evasion, and virulence.^2,3^ A distinguishing feature of this VOC is the large number of residue changes detected in the Spike protein (hereafter Spike) (~30) versus all previously characterized viral strains. Of particular concern are the numerous (~15) changes in the receptor-binding domain (RBD) of its Spike (**Fig. 1A**), suggesting the possibility that they may either dampen or evade individuals’ immune responses generated by vaccination and/or previous infection. Immune evasion may have serious consequences, potentially leading to increased incidence of infection, reinfection and/or further enhancing viral fitness during evolution in the event of uncontrolled spread around the globe. Early data suggest substantial but not complete immune evasion.^4^ A second concern is that therapeutic proteins, such as monoclonal antibody cocktails or nanobodies, developed against previously characterized SARS-CoV-2 variants may no longer be effective in neutralizing the Omicron VOC, ultimately leading to increased morbidity and mortality.

**Figure 1:**
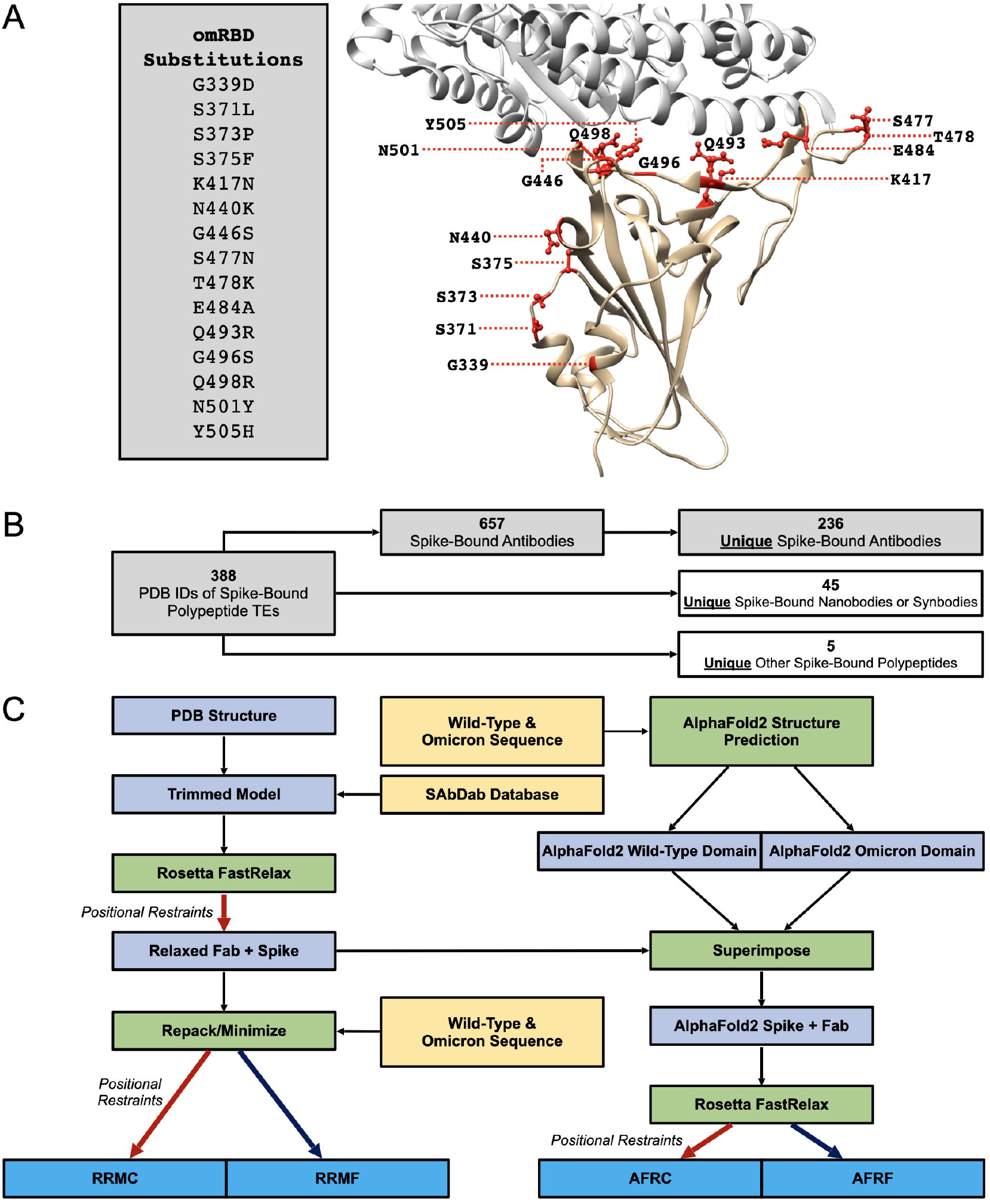
**(A)** Overview of residues of Spike of SARS-CoV-2 that are substituted in the Omicron VOC receptor-binding domain (omRBD) (tan). RBD contains the receptorbinding motif (RBM) which interacts with ACE2 (light gray) (PDB ID: 6M0J). **(B)** Overall breakdown of all 388 PDB IDs containing Spike-bound polypeptide therapeutic entities (TEs) (as of November 27, 2021). This study focuses on antibodies. **(C)** Workflow to generate predicted structures of omRBD in complex with various polypeptide TEs, including antibodies, based on experimentally solved structures.

At the core of concerns regarding immune evasion is molecular recognition in three dimensions (3D) between the Spike and viral ligands, including at least one of its established cellular receptors (ACE2), neutralizing antibodies, therapeutic antibodies, and other binding proteins. Several fundamental queries must be urgently addressed. First, is the binding of these agents to the Omicron VOC Spike substantially altered compared to previous variants of SARS-CoV-2 (*e.g*., the Delta VOC, or the original Wuhan-Hu-1 strain)? Moreover, do the numerous sequence changes accrued in the Omicron VOC Spike alter its shape? If so, do these changes lead to a meaningful remodeling of interfaces formed with various binding proteins? Finally, are these remodeled interfaces likely to weaken the recognition of Omicron VOC Spike by some or all of the neutralizing entities and do these molecular changes correlate with neutralization data, which will become available in the near future? Experimental answers to each of these currently open questions would likely have profound impact in biology, medicine, and public health policy in an ongoing global airborne pandemic.

To begin to shed light on these questions at the atomic level in 3D and aid experimental characterization and potential redesign of therapeutic entities (TEs), we turned to the wealth of experimental structures of (previous variants) Spike-TE interactions generated by the structural biology community and freely available from the Protein Data Bank (PDB),^5^ and recent advances in artificial intelligence/machine learning (AI/ML)-based protein structure prediction.^6–8^ These resources can be critical in understanding and even optimizing the immune response against SARS-CoV-2 variants; for example, our group utilized a structure-centered approach to initially identify an immunogenic epitope that remains universal across all major emergent variants.^9,10^ Focusing our attention on the receptor-binding domain (RBD) of the Spike, we generated computed structural models of the Omicron VOC RBD (omRBD) in the unbound and bound states with all polypeptide TEs for which there are atomic level structures archived in the PDB (388 multi-protein complex structures, some of which contain more than one distinct polypeptide, for a total of 286 unique bound polypeptides). Alterations in individual Spike-TE interfaces and a large-scale analysis of these complexes may provide insights into the molecular bases for dampened immune response and suggest avenues for redesign of TEs (*e.g*., antibody cocktails) as effective countermeasures against the Omicron VOC.

While our analyses still are ongoing and evolving, we believe that making these models widely available will aid wet laboratory researchers who are studying Omicron VOC Spike-antibody recognition in generating hypotheses for altered recognition, and enable further characterization of complexes via more computationally-expensive methods. We strongly caution that antibody-binding affinity, while important, is but one of many other factors involved in immune evasion or any complex biological response. Advanced structure prediction tools such as AlphaFold2 (AF2)^8^ and RoseTTAFold^6^ although very powerful, have not been trained to predict changes in binding affinity. In many computational affinity prediction approaches, there is a tradeoff between computational speed and prediction accuracy. Therefore, overly hasty and caveat-free application of modeling pipelines for predicting antibody binding affinities along with the application of calculated binding energies to predict viral neutralization is decidedly unwise and likely misguided. Given that SARS-CoV-2 Spike recognition is central to biochemical interactions between our immune systems and the virus, and the urgent unmet need to understand the efficacy of immune response against the Omicron VOC, we believe that structure-based and large-scale analyses of Spike interfaces will provide key, albeit qualitative, insights. We are, therefore, sharing these models with the scientific and medical research communities writ large. As such, all of the computed structural models together with associated summary sheets described in this work are available online (https://github.com/sagark101/omicron_models). This report includes an initial set of 38 complexes of 20 binding entities; furthermore, we will be continually updating the computed models and our analyses on the GitHub repository as additional data become available.

## Results

### Curation of SARS-CoV-2 Spike Structures in the PDB

We have curated all structures of Spike bound to therapeutic polypeptide entities (388 distinct PDB IDs as of November 27, 2021) (**Fig. 1B**). The cocktail status of PDB IDs was determined (cocktail was defined as two or more unique bound proteins in a single multi-protein complex). Bound proteins/peptides were listed for each PDB ID and, if a cocktail, specifically by polypeptide chain(s). All Spike-bound therapeutic entities were subdivided into three possible groups: antibody, nanobody/synbody, and other (**Table 1**).

**Table 1:**
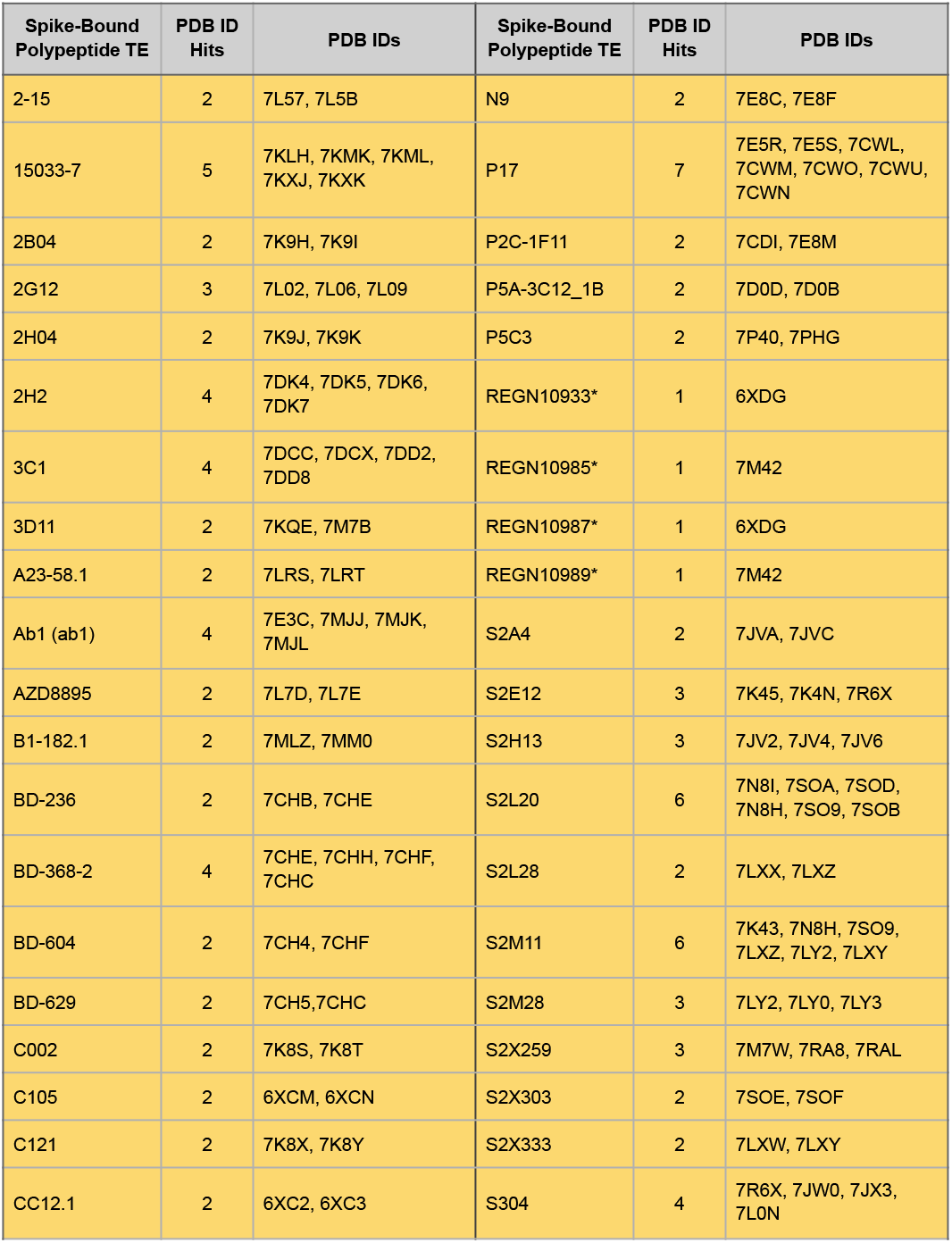
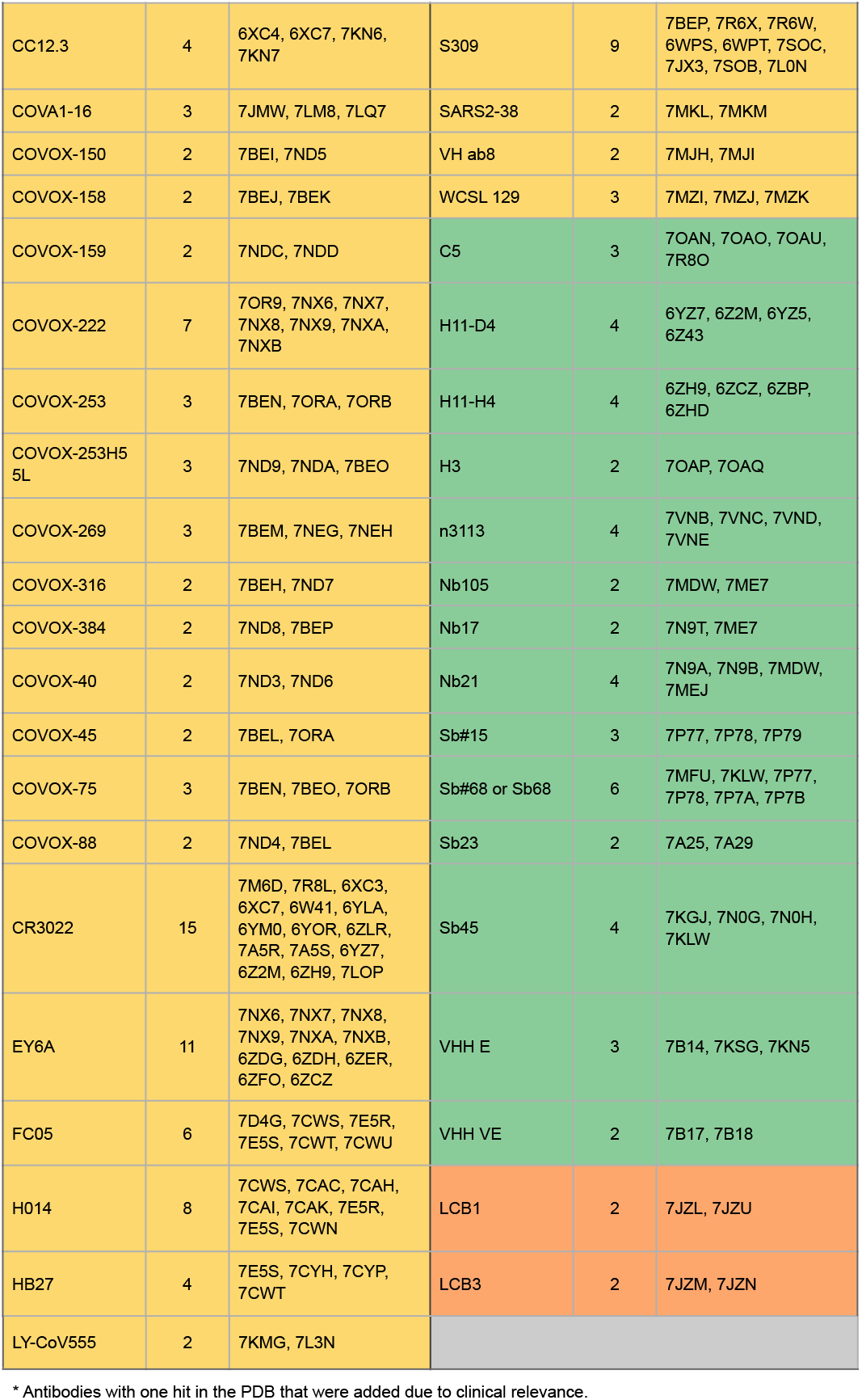
Summary of Spike-bound polypeptide TEs occurring in at least two PDB IDs. Color indicates subdivision of bound protein: antibody (yellow), nanobody/synbody (green), and other (orange).

| Spike-Bound Polypeptide TE | PDB ID Hits | PDB IDs | Spike-Bound Polypeptide TE | PDB ID Hits | PDB IDs |
| --- | --- | --- | --- | --- | --- |
| 2-15 | 2 | 7L57, 7L5B | N9 | 2 | 7E8C, 7E8F |
| 15033-7 | 5 | 7KLH, 7KMK, 7KML, 7KXJ, 7KXK | P17 | 7 | 7E5R, 7E5S, 7CWL, 7CWM, 7CWO, 7CWU, 7CWN |
| 2B04 | 2 | 7K9H, 7K9I | P2C-1F11 | 2 | 7CDI, 7E8M |
| 2G12 | 3 | 7L02, 7L06, 7L09 | P5A-3C12_1B | 2 | 7D0D, 7D0B |
| 2H04 | 2 | 7K9J, 7K9K | P5C3 | 2 | 7P40, 7PHG |
| 2H2 | 4 | 7DK4, 7DK5, 7DK6, 7DK7 | REGN10933* | 1 | 6XDG |
| 3C1 | 4 | 7DCC, 7DCX, 7DD2, 7DD8 | REGN10985* | 1 | 7M42 |
| 3D11 | 2 | 7KQE, 7M7B | REGN10987* | 1 | 6XDG |
| A23-58.1 | 2 | 7LRS, 7LRT | REGN10989* | 1 | 7M42 |
| Ab1 (ab1) | 4 | 7E3C, 7MJJ, 7MJK, 7MJL | S2A4 | 2 | 7JVA, 7JVC |
| AZD8895 | 2 | 7L7D, 7L7E | S2E12 | 3 | 7K45, 7K4N, 7R6X |
| B1-182.1 | 2 | 7MLZ, 7MM0 | S2H13 | 3 | 7JV2, 7JV4, 7JV6 |
| BD-236 | 2 | 7CHB, 7CHE | S2L20 | 6 | 7N8I, 7SOA, 7SOD, 7N8H, 7SO9, 7SOB |
| BD-368-2 | 4 | 7CHE, 7CHH, 7CHF, 7CHC | S2L28 | 2 | 7LXX, 7LXZ |
| BD-604 | 2 | 7CH4, 7CHF | S2M11 | 6 | 7K43, 7N8H, 7SO9, 7LXZ, 7LY2, 7LXY |
| BD-629 | 2 | 7CH5, 7CHC | S2M28 | 3 | 7LY2, 7LY0, 7LY3 |
| C002 | 2 | 7K8S, 7K8T | S2X259 | 3 | 7M7W, 7RA8, 7RAL |
| C105 | 2 | 6XCM, 6XCN | S2X303 | 2 | 7SOE, 7SOF |
| C121 | 2 | 7K8X, 7K8Y | S2X333 | 2 | 7LXW, 7LXY |
| CC12.1 | 2 | 6XC2, 6XC3 | S304 | 4 | 7R6X, 7JW0, 7JX3, 7L0N |
| CC12.3 | 4 | 6XC4, 6XC7, 7KN6, 7KN7 | S309 | 9 | 7BEP, 7R6X, 7R6W, 6WPS, 6WPT, 7SOC, 7JX3, 7SOB, 7L0N |
| COVA1-16 | 3 | 7JMW, 7LM8, 7LQ7 | SARS2-38 | 2 | 7MKL, 7MKM |
| COVOX-150 | 2 | 7BEI, 7ND5 | VH ab8 | 2 | 7MJH, 7MJI |
| COVOX-158 | 2 | 7BEJ, 7BEK | WCSL 129 | 3 | 7MZI, 7MZJ, 7MZK |
| COVOX-159 | 2 | 7NDC, 7NDD | C5 | 3 | 7OAN, 7OAO, 7OAU, 7R8O |
| COVOX-222 | 7 | 7OR9, 7NX6, 7NX7, 7NX8, 7NX9, 7NXA, 7NXB | H11-D4 | 4 | 6YZ7, 6Z2M, 6YZ5, 6Z43 |
| COVOX-253 | 3 | 7BEN, 7ORA, 7ORB | H11-H4 | 4 | 6ZH9, 6ZCZ, 6ZBP, 6ZHD |
| COVOX-253H5 5L | 3 | 7ND9, 7NDA, 7BEO | H3 | 2 | 7OAP, 7OAQ |
| COVOX-269 | 3 | 7BEM, 7NEG, 7NEH | n3113 | 4 | 7VNB, 7VNC, 7VND, 7VNE |
| COVOX-316 | 2 | 7BEH, 7ND7 | Nb105 | 2 | 7MDW, 7ME7 |
| COVOX-384 | 2 | 7ND8, 7BEP | Nb17 | 2 | 7N9T, 7ME7 |
| COVOX-40 | 2 | 7ND3, 7ND6 | Nb21 | 4 | 7N9A, 7N9B, 7MDW, 7MEJ |
| COVOX-45 | 2 | 7BEL, 7ORA | Sb#15 | 3 | 7P77, 7P78, 7P79 |
| COVOX-75 | 3 | 7BEN, 7BEO, 7ORB | Sb#68 or Sb68 | 6 | 7MFU, 7KLW, 7P77, 7P78, 7P7A, 7P7B |
| COVOX-88 | 2 | 7ND4, 7BEL | Sb23 | 2 | 7A25, 7A29 |
| CR3022 | 15 | 7M6D, 7R8L, 6XC3, 6XC7, 6W41, 6YLA, 6YM0, 6YOR, 6ZLR, 7A5R, 7A5S, 6YZ7, 6Z2M, 6ZH9, 7LOP | Sb45 | 4 | 7KGJ, 7N0G, 7N0H, 7KLW |
| EY6A | 11 | 7NX6, 7NX7, 7NX8, 7NX9, 7NXA, 7NXB, 6ZDG, 6ZDH, 6ZER, 6ZFO, 6ZCZ | VHH E | 3 | 7B14, 7KSG, 7KN5 |
| FC05 | 6 | 7D4G, 7CWS, 7E5R, 7E5S, 7CWT, 7CWU | VHH VE | 2 | 7B17, 7B18 |
| H014 | 8 | 7CWS, 7CAC, 7CAH, 7CAI, 7CAK, 7E5R, 7E5S, 7CWN | LCB1 | 2 | 7JZL, 7JZU |
| HB27 | 4 | 7E5S, 7CYH, 7CYP, 7CWT | LCB3 | 2 | 7JZM, 7JZN |

|  |  |  |
| --- | --- | --- |
| LY-CoV555 | 2 | 7KMG, 7L3N |

\* Antibodies with one hit in the PDB that were added due to clinical relevance.

From the pool of antibodies with at least two exemplars (plus the REGN antibodies) in the PDB, we selected RBD-binding antibodies across all four Barnes Classes (numbered 1-4) for modeling in this initial report.^11^ Eventually, we anticipate performing similar analyses on all proteins in **Table 1** and more as they accumulate in the PDB. For the current study, we pursued antibodies that definitively belong to a particular Barnes Class based on available literature. Supervised selection criteria for PDB ID for a given antibody were prioritized as follows: (i) higher-resolution, (ii) whether residues in and near the Spike-binding site were resolved, (iii) absence of Spike substitutions (*i.e*., feature Wuhan-Hu-1 amino acid sequence), and (iv) solely contained our intended antibody (as opposed to a cocktail). In cases where a PDB ID contained a cocktail bound to Spike, the additional antibodies were also included for modeling. We have ultimately selected 20 of 236 unique Spike-bound antibodies available from the PDB as representative cases of RBD-binding antibodies for detailed analysis in the context of the Omicron VOC (**Table 2**).

**Table 2:** Representative RBD-bound antibodies across Barnes Classes 1-4 selected for initial structural modeling and analysis.

| RBD-Bound Antibody | Barnes Class | PDB ID | Study Modeling Code |
| --- | --- | --- | --- |
| A23-58.1 | 1 | 7LRT | 7LRT_1, 7LRT_2, 7LRT_3 |
| B1-182.1 | 1 | 7MM0 | 7MM0_1 |
| C105 | 1 | 6XCM | 6XCM_1, 6XCM_2 |
| CC12.3 | 1 | 6XC4 | 6XC4_1*, 6XC4_2 |
| COVOX-222 | 1 | 7NX6 | 7NX6_2* |
| COVOX-253 | 1 | 7ORA | 7ORA_1, 7ORA_4 |
| REGN10933 | 1 | 6XDG | 6XDG_1* |
| S2X259 | 1 | 7M7W | 7M7W_2, 7M7W_4 |
| C002 | 2 | 7K8S | 7K8S_1, 7K8S_2, 7K8S_3* |
| C121 | 2 | 7K8X | 7K8X_1, 7K8X_2 |
| LY-CoV555 | 2 | 7KMG | 7KMG_1, 7KMG_2 |
| REGN10987 | 3 | 6XDG | 6XDG_2* |
| S309 | 3 | 7BEP, 7R6X | 7BEP_1*, 7BEP_4, 7R6X_1 |
| CR3022 | 4 | 6YLA | 6YLA_1*, 6YLA_2 |
| EY6A | 4 | 6ZDH, 7NX6 | 6ZDH_1, 6ZDH_2, 6ZDH_3, 7NX6_1 |
| S304 | 4 | 7R6X | 7R6X_2 |

\* Modeled complexes presented in Fig. 4-9.
Note: Additional antibodies were included in this initial subset for modeling because they are present in cocktail complexes: COVOX-45 (PDB ID: 7ORA), COVOX-384 (PDB ID: 7BEP), S2E12 (PDB ID: 7R6X), and S2H97 (PDB ID: 7M7W).

### AF2 Modeling of Omicron Spike and Fragments

As no experimentally-determined structures of Omicron VOC proteins are currently available, we used AF2 to generate models of omRBD and various Omicron VOC Spike fragments (S1 domain, full monomer, full trimer). We modeled full Omicron VOC Spike sequences as monomers in AF2 and attempted to model trimers as well. Predicted structures of individual domains and the Spike monomer are deemed credible. The pLDDT scores are generally higher than 75 for the top models of monomers and more than 85 for the top models of the individual domains. A score of above 70 for a predicted structure is considered to be in the confident range.^8^ The AF2 algorithm predicts five structural models and ranks them according to their predicted local-distance difference (pLDDT) score,^8,12^ which is reported in the 0-100 range. Confidence in predicted structure is deemed to be high when the score is more than 90, medium when the score is between 70 and 90, low when the score is less than 70, and very low when the score is less than 50. Pairwise Root-Mean-Square-Deviation (RMSD) values comparing Cα atomic positions for experimental structures for several other variant Spike fall in the range of 0.3-1.3 Å (**Supplementary Table 1**). We have further benchmarked the AF2-predicted monomers by comparison with several other variant Spike experimental structures in the PDB, including Alpha, Beta, Gamma, Delta, and P1 variants, and a variant RBD produced by directed evolution.^13^ Cα RMSD values range from 0.4-0.8 Å (**Supplementary Table 2**). Initial attempts at AF2 computational modeling of the trimeric full Spike structure were unsuccessful, producing one reasonable monomeric chain and two misfolded chains (data not shown).

AF2-predicted single-domain models of the omRBD are also similar to Rosetta-minimized PDB-based structures of omRBD bound to various binding entities. Cα RMSD values range from 0.5-2.2 Å (**Supplementary Table 3**), which is comparable to the structural variation observed within the set of experimental Spike structures. Domain N- and C-termini are typically less well-structured in both experimental and predicted models, and, therefore, represent a substantial source of the structural deviations. Correspondingly, according to the predicted Template Model (pTM) scores that AF2 assigns to each residue (**Fig. 2, Fig. S1**), there is high confidence in the portion of the domain that interacts with ACE2, and lower confidence closer to both termini. This finding is consistent with the known flexibility of the Spike in this region, which allows the RBD to hinge between open and closed conformations. The greatest structure prediction uncertainty within the RBD polypeptide chain occurs in the α-helix composed of residues 364-376, in which three sites are substituted in the Omicron VOC (**Fig. 2, inset**). Notably, among these substitutions is S373P, which would be expected to cause disruption or kinking of the α-helix compared to wild-type. It should also be noted that these residues also have more positive (*i.e*., unfavorable) scores in the Rosetta-computed models, indicating a similar lack of confidence in the correct folding around those substituted residues. Finally, when assessing binding involving the regions with low-confidence predictions, one must keep in mind that the low confidence of the unbound structure propagates into lower confidence in the energetic calculations at those sites.

**Figure 2:**
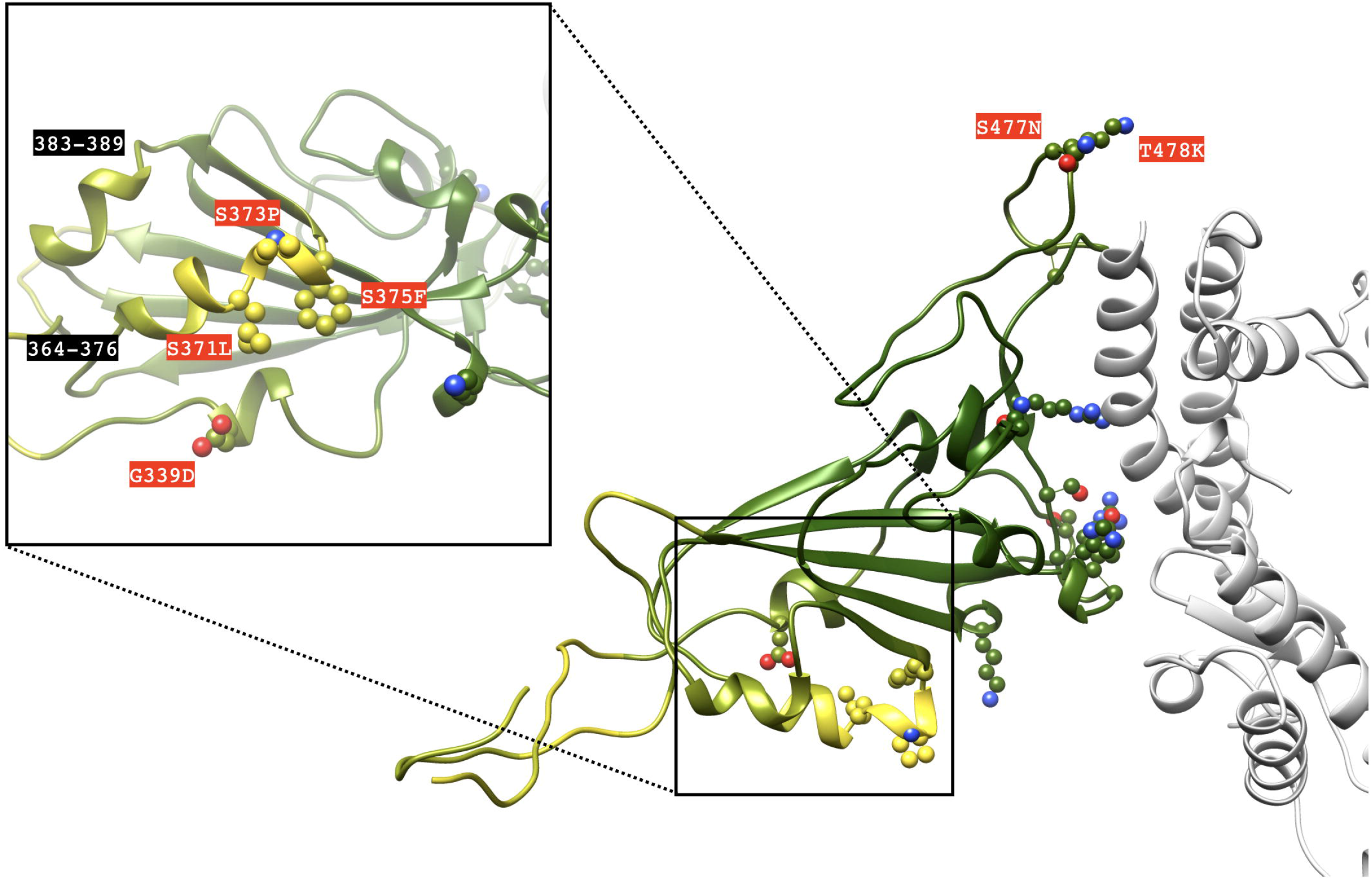
AFRC model of omRBD in complex with ACE2 (light gray) (PDB ID: 6M0J). omRBD residues are colored by AF2-reported pTM in the generated model (yellow-to-green spectrum corresponds to low-to-high confidence). Inset: substituted residues with low pTM in comparison to other substitutions are highlighted (red labeling).

### Overview of Variant Modeling Approach

We have previously developed an “*in silico* mutagenesis” and analysis pipeline for studying the ongoing populational co-evolution of SARS-CoV-2.^14^ In our pipeline, amino acid substitutions observed in sequenced SARS-CoV-2 genomes obtained from the GISAID database^15,16^ are introduced in available crystal structures or computed structural models of proteins and their energetic impact is qualitatively evaluated. We have previously focused on evaluating how observed mutations affect the structure and stability of individual proteins and domains for SARS-CoV-2.^14^ We adapted this pipeline (**Fig. 1C**) to the evaluation of omRBD in complex with ACE2 and bound antibodies, etc.

### Complex Model Generation

To increase computational efficiency during modeling, structures were truncated to include only RBD and binding regions of the modeled binding partners. Antibody binding regions were identified with reference to the Structural Antibody Database (SAbDAb).^17^ As no experimentally determined structures of omRBD are available, we used two starting points for generating omRBD models: (i) RBD atomic coordinates available in X-ray structures archived in the PDB, and (ii) AF2 models of the free RBD, which were superimposed on the RBD atomic coordinates in PDB structures. For each type of starting structure, we applied two conformational sampling approaches to obtain Rosetta energy-minimized models of the complexes. Interface side-chain rotamers were repacked and gradient-based minimization of the Rosetta-calculated energy was performed with and without positional coordinate restraints on the protein backbone atoms, leading to four models for each modeled complex with a given variant. We hereafter refer to these models as follows: RRMC, Rosetta Repack Minimize Constrained; RRMF, Rosetta Repack Minimize Free; AFRC, AF2 Repack-minimize Constrained; and AFRF, AF2 Repack-minimize Free. Repack-Minimize is performed using the Rosetta energy function in all cases.^18^ Having models generated with various degrees of conformational constraints during optimization is intended to provide representative low-energy conformations from the native ensemble of the complex as our conformational sampling approach involves a tradeoff between computational speed and extensive evaluation of the native ensemble.

### Consensus Scoring

With eight models (four for omRBD, four for Wuhan-Hu-1 RBD) for each binding partner in hand, we next examined these models for altered interaction patterns and evaluated per-residue energies of interaction to identify the consensus effect of amino acid substitutions. Examination of energy changes across all four types of models serves as a measure of confidence in the identified interaction patterns. For each model, we have also used a threshold value of 1.4 Rosetta Energy Units (REU) to identify changes in residue energies. For each residue, one “+” is assigned per model in which the interface destabilization energy due to substitution exceeds +1.4 REU, and one “-” is assigned per model in which it is below −1.4 REU, with + and - canceling each other, and neither is assigned for models where the absolute value of interface energy change was below 1.4 REU. For example, a particular mutation is scored as ++++ (high-consensus destabilizing) if all four models involve calculated interface energy increases larger than 1.4 REU. A score of ++ was assigned if two of the four models feature destabilization greater than the threshold value and two others do not have significant energy changes with absolute value greater than 1.4 REU, or if three are above 1.4 REU and one is below −1.4 REU. A “*” is assigned to a substitution when two or more models predict conflicting but substantial changes, indicating low consensus in our prediction. Of the 38 antibody complexes presented in this initial report, 91 substitutions exceeded the absolute value of 1.4 REU to be noteworthy. Of those, 13 were ++++ (14%) and 11 were +++ (12%), whereas only five were --- (6%), and three were ---- (3%). These trends indicate that several key interactions predicted with high consensus are likely disruptive, but there are a small number of cases in which we can anticipate some compensation by a second substitution. As cautionary notes, these observations reflect the consequences to substituted residues only, and may or may not directly capture effects in the surrounding residues; these effects may also contribute to the changes in overall binding energy upon substitution to some extent. More information can be found in the **Supplementary Materials**.

## Results

### Analysis of ACE2 Binding Interface

To investigate the impact of omRBD residue substitutions on ACE2 binding, we examined the modeled complexes of omRBD-ACE2 and compared them to the experimental structure of the original Wuhan-Hu-1 RBD-ACE2 complex (**Fig. 3**). All of the amino acid substitutions at the ACE2 interface are accommodated in a favorable geometry and appear to make several favorable intra- and inter-molecular contacts consistent with binding to ACE2 being preserved despite the large number of amino changes observed in the RBD. Many of the substitutions or sites of substitutions in omRBD were observed individually in other variants (N501Y, first seen in Alpha and Gamma; K417N and E484K, first seen in Beta; T478K; first seen in Delta). Some of these substitutions have been observed in pairwise combinations in RBD domains evolved for increasing affinity to the ACE2 receptor.^13^ Analyses of synonymous and non-synonymous mutations in the gene encoding the Omicron VOC Spike compared to previously detected SARS-CoV-2 variants indicate that several key sites of substitution have likely undergone positive selection, suggesting that the observed amino acid substitutions, particularly those occurring in the receptor-binding motif, are beneficial to the virus.^19^ One key question for a markedly altered VOC such as Omicron is whether these individual favorable substitutions will interact favorably (positive cooperativity/epistasis) or unfavorably (negative cooperativity/epistasis), and what might be the structural basis for such an effect? Our computed structural models document that for some omRBD substitutions that cluster in a single binding region, positive cooperativity may arise as a result of an omRBD-specific extensive interdigitated largely polar interaction network (featuring Y501, R498, S496, R493 from the RBD; *K353, D38, E35* from ACE2: italics indicate ACE2 residue numbering) between the omRBD substitutions and ACE2 residues in the binding site (**Fig. 3**). This enthalpically-favorable interaction network is markedly smaller in the original RBD where the interface features individual residue pairs (Q493-*E35*) or small clusters (Q498-*D38*-Y449) of hydrogen bonds (**Fig. 3**). In the extensive network of predicted interactions, the omRBD-ACE2 interface bears some resemblance to other evolved protein-protein interfaces with optimized, often interdigitated sidechain-sidechain interactions.^20,21^ Analysis of calculated energy changes (**Fig. 3C**) suggests that three substitutions contribute favorably to the binding (with high consensus) while K417N leads to the loss of an interface hydrogen bond. However, N417 is involved in two intramolecular hydrogen bonds that may increase the stability of the bound conformation. Overall, the models reported here suggest that ACE2 binding will be robust with the emergence of a multi-residue network of interactions and likely enhanced compared to the Wuhan-Hu-1 RBD.

**Figure 3:**
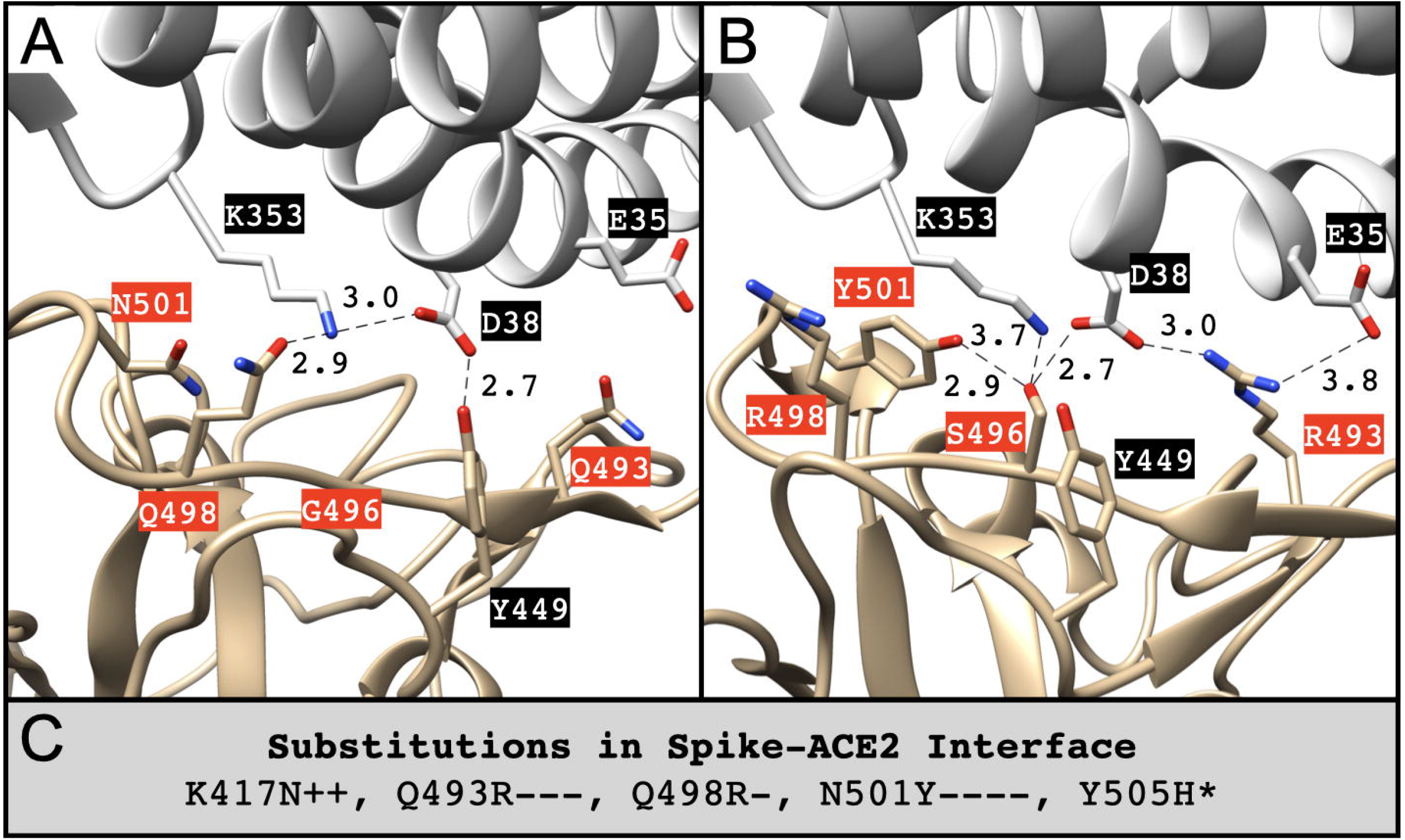
Key substituted residues (labeled in red) at the interface between Spike (tan) and ACE2 (dim gray) that undergo notable energy changes in the Omicron VOC based on its **(A, B)** AFRF model (PDB ID: 6M0J). Length unit of non-covalent bonds (dotted lines) between Spike and ACE2 is Å. **(C)** Substituted residues of Spike involved in the binding interface and their relative energy changes across four predicted models (RRMC, RRMF, AFRC, AFRF). “+” indicates destabilization and “-” indicates stabilization (absolute magnitude larger than 1.4 REU). Number of “+” or “-” symbols indicates our confidence in the prediction (three or four: high, two: moderate, one: low), * indicates a situation where there are conflicting predictions from two or more methods.

### Analysis of Antibody Binding Interfaces

One of the more plausible scenarios advanced to explain the origin of the Omicron VOC is long-term circulation in an immunocompromised subject (presumably human) with persistent SARS-CoV-2 infection.^22^ Within such an environmental niche, it can be reasonably posited that omRBD has accrued substitutions endowing it with both increased infectivity and immune escape properties. Indeed, several sites of substitutions in the Omicron VOC have been observed in other variants that exhibit dampened antibody binding^23^ and in yeast-based deep mutational escape mapping experiments carried out with a large set of antibodies.^24–26^ These previous results suggest that the Omicron VOC might be capable of immune evasion from neutralizing and therapeutic antibodies.^27^ As this VOC appears to have many escape-endowing substitutions, a key question is whether their individual effects will be additive or display cooperativity (positive or negative). In other words, can the weakened affinity due to one substitution be compensated by additional interactions due to another substitution? Because these interactions are inherently 3D structure-mediated phenomena, we reasoned that computed structural models of omRBD-antibody complexes may provide insights into possible interactions between effects of individual substitutions.

### Antibodies Targeting SARS-CoV-2

SARS-CoV-2 antibodies have been divided by Barnes, Björkman, and colleagues into four classes (Barnes Classes), depending on whether their binding sites overlap with the ACE2-binding site and the orientation of RBD for accessibility (up or down).^11^ We have computed structural models of omRBD bound to representative antibodies from each Barnes Class, including three therapeutic antibodies (REGEN-COV cocktail containing two Regeneron Pharmaceuticals antibodies, and S309 which has been adapted by Vir Biotechnology and GlaxoSmithKline), and analyzed their interfaces in detail.

#### Class 1 Antibodies

Class 1 neutralizing antibodies have epitopes on the Spike that overlap with the ACE2-binding site and can only bind to RBD in the up (open) conformation.^11^ These antibodies are typically defined by their short CDRH3 loops. Various examples include C102, C105, B38, CC12.3, COVOX-222, COVOX-253.^11^ We opted to analyze models of Spike-antibody complexes for CC12.3 and COVOX-222 and evaluate the impact of omRBD-associated substitutions.

CC12.3 is a VH3-53 public-domain antibody that is elicited from infection with COVID-19.^28^ We investigated models of the omRBD-CC12.3 complex to assess the impact of interfacial substitutions (**Fig. 4**). Notably, but as expected, the consensus among three out of four models is that K417N will evoke a destabilizing decrease in binding energy (consistent with the aforementioned antibody studies), likely due to the loss of two interactions made by K417: a salt bridge and hydrogen bond with CC12.3 (**Fig. 4A, 4B**). N417 instead makes intramolecular hydrogen bonding interactions. This finding is consistent with other studies that document at least 100-fold reduction in antibody binding as a result of K417N.^23,25,29^ The other two substituted residues (N501Y and Y505H) at the interface have a moderate impact (low consensus) on CC12.3 binding to the Omicron VOC Spike (**Fig. 4C-4E**). Interestingly, N501Y has been documented to have a negligible impact (0.1-fold reduction) on CC12.3 binding.^23,25^

**Figure 4:**
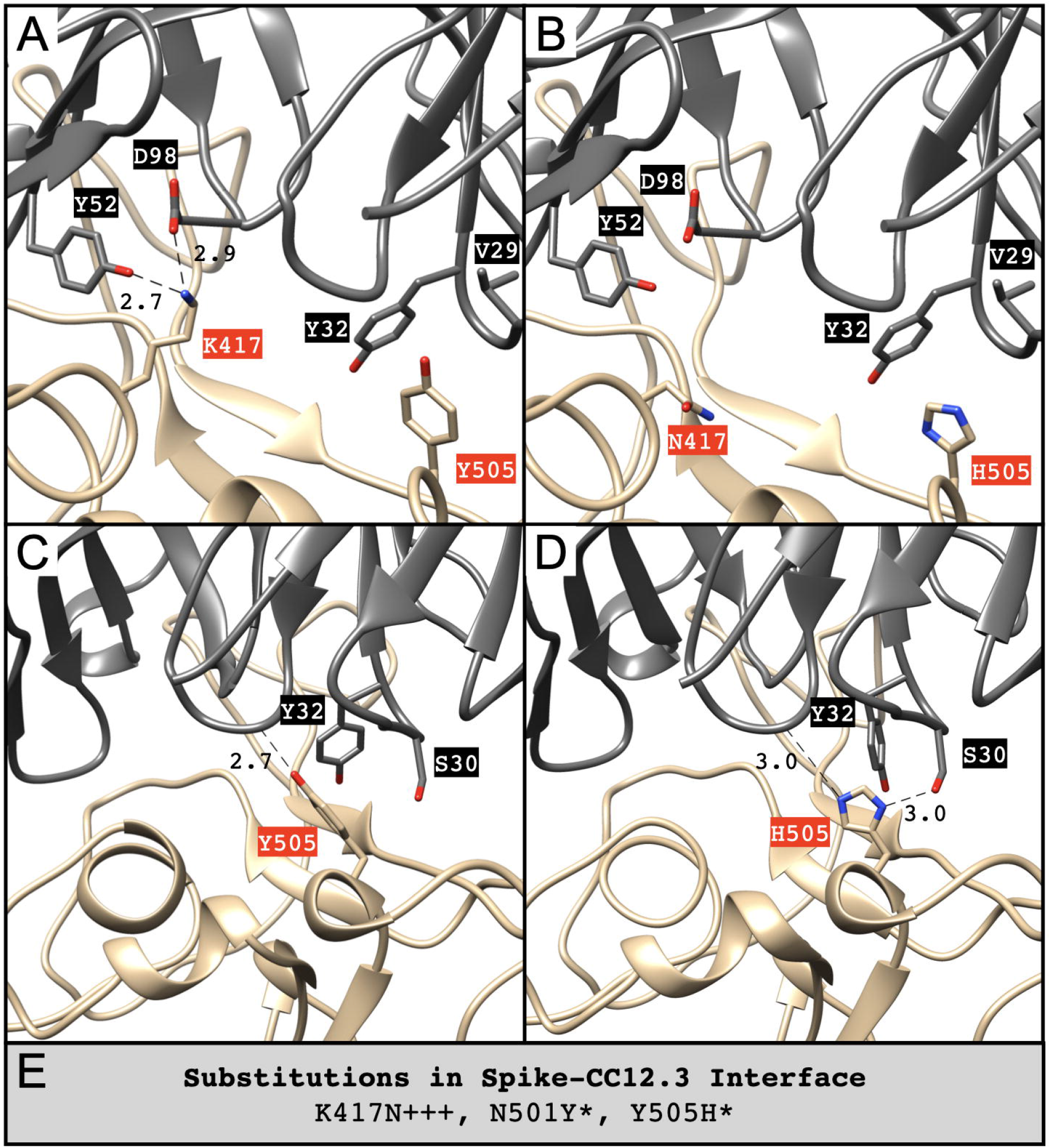
Key substituted residues (red) in the interface between Spike (tan) and CC12.3 (dim gray) that undergo notable energy changes in the Omicron VOC based on its **(A, B)** RRMC and **(C, D)** AFRF models (PDB ID: 6XC4). Length unit of non-covalent bonds (dotted lines) between Spike and antibody is Å. **(E)** Substituted residues of Spike involved in the binding interface and their relative energy changes across four predicted models (RRMC, RRMF, AFRC, AFRF). “+” indicates destabilization and “-” indicates stabilization (absolute magnitude larger than 1.4 REU). Number of “+” or “-” symbols indicates our confidence in the prediction (three or four: high, two: moderate, one: low), * indicates a situation where there are conflicting predictions from two or more methods.

COVOX-222 (also referred to as Fab 222 or mAb 222) is a Class 1 neutralizing antibody which also belongs to the VH3-53 public domain.^30,31^ Most of our models indicate that K417N is destabilizing, and one model finds that Q493R is stabilizing. K417N results in the loss of one salt bridge and two hydrogen bonds (**Fig. 5A,5B**). Conversely, Q493R permits the formation of two backbone hydrogen bonds (**Fig. 5C,5D**). All four of our models of omRDB-COVOX-222 indicate that Y505H is destabilizing at the interface (**Fig. 5**), potentially due to the loss of a hydrogen bond (**Fig. 5C,5D**). Other interfacial substitutions that may have a modest favorable effect (low consensus) on binding are G496S, Q498R, and (moderate consensus) N501Y (**Fig. 5E**).

**Figure 5:**
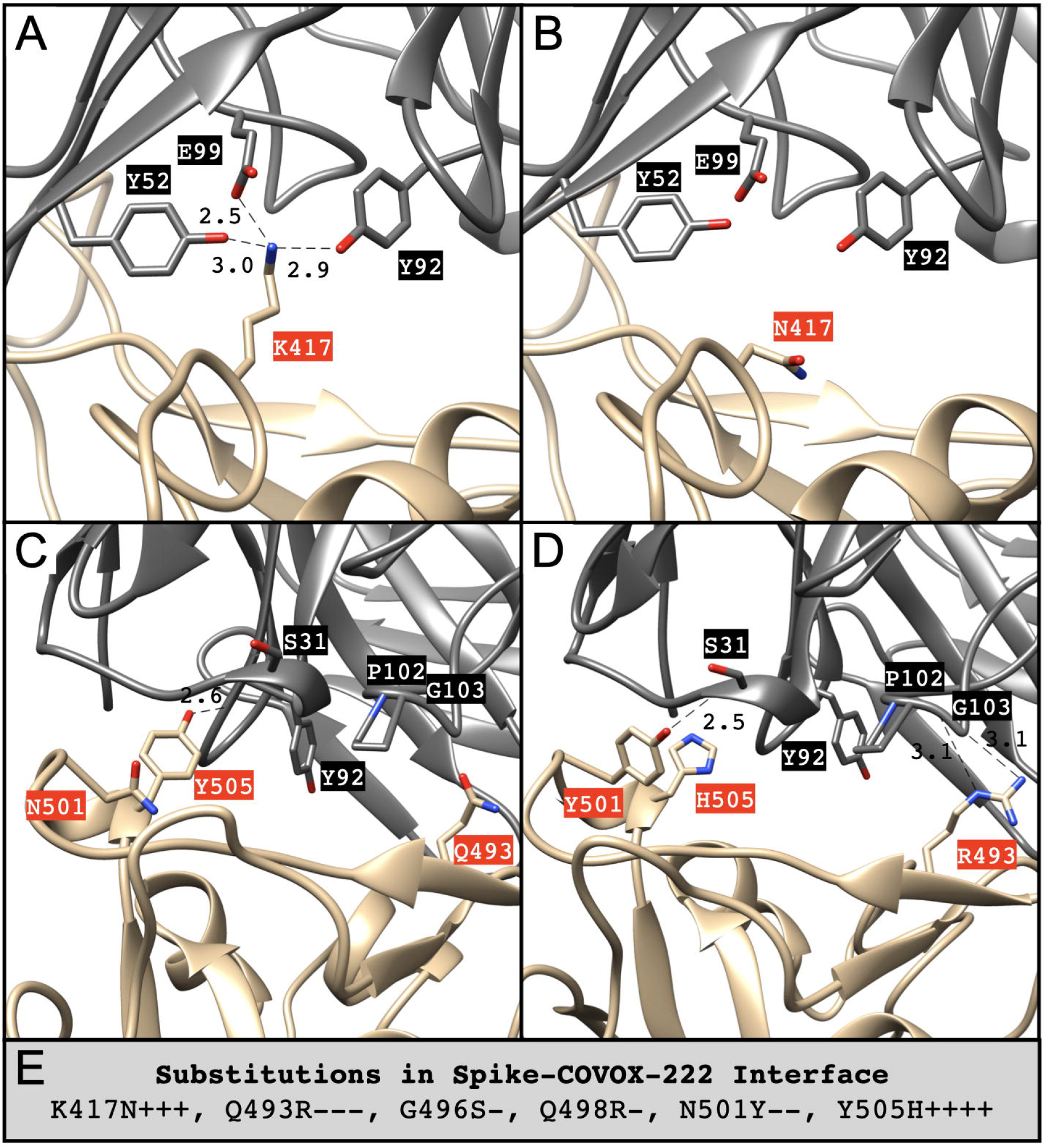
Key substituted residues (red) in the interface between Spike (tan) and COVOX-222 (dim gray) that undergo notable energy changes in the Omicron VOC based on its **(A, B)** RCRF and **(C, D)** AFRC models (PDB ID: 7NX6). Length unit of non-covalent bonds (dotted lines) between Spike and antibody is Å. **(E)** Substituted residues of Spike involved in the binding interface and their relative energy changes across four predicted models (RRMC, RRMF, AFRC, AFRF). “+” indicates destabilization and “-” indicates stabilization (absolute magnitude larger than 1.4 REU). Number of “+” or “-” symbols indicates our confidence in the prediction (three or four: high, two: moderate, one: low), * indicates a situation where there are conflicting predictions from two or more methods.

#### Class 2 Antibodies

Class 2 antibodies are characterized by Spike epitopes that overlap the ACE2-binding site and can bind RBD irrespective of its conformation (up/open or down/closed).^11^ In comparison to Class 1, these antibodies tend to have larger CDRH3 loops and bridge RBDs. Examples include C002, C104, C119, C121, C144, COVA2-39, 5A6, P2B-2F6, Ab2-4, and BD23.^11^ We have assessed computed structural models of the omRBD-C002 complex as a representative case.

C002 is a neutralizing antibody originally isolated from COVID-19-convalescent sera^32^ and encoded by VH3-30 and VK1-39.^11^ All four models of omRBD-C002 highlight the markedly destabilizing effects of K417N and E484A at the interface (**Fig. 6**). K417N is predicted to lose a salt bridge with C002 (**Fig. 6A,6B**) while E484A loses both a salt bridge and hydrogen bond (**Fig. 6C,6D**). Furthermore, three of four models highlight the energetic unfavorability of Q493R for binding, which may result in the loss of two hydrogen bonds. Other potential consequences from the Omicron VOC include the stabilization by G496S according to one model due to the formation of novel sidechain interactions (**Fig. 6E,6F**). Our findings are consistent with a previous study^33^ which discovered that Omicron-relevant substituted residues retain high escape fraction values, specifically for E484A (1.0) and Q493R (0.979).^23^ Overall, predicted loss of interactions at four sites of omRBD suggests that binding to C002 may be substantially compromised.

**Figure 6:**
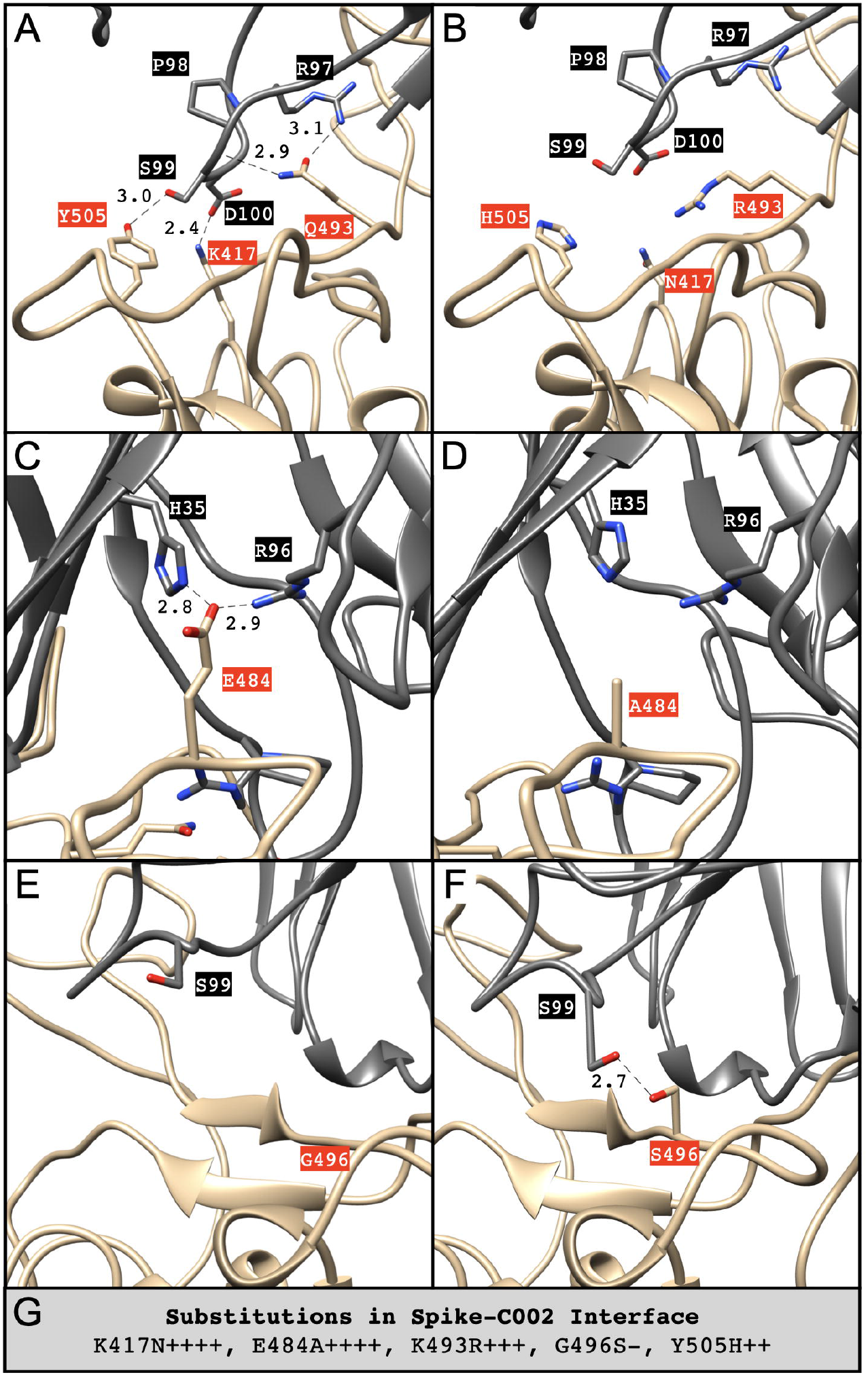
Key substituted residues (red) in the interface between Spike (tan) and C002 (dim gray) that undergo notable energy changes in the Omicron VOC based on its **(A, B, C, D)** RRMC and **(E, F)** AFRC models (PDB ID: 7K8S). Length unit of non-covalent bonds (dotted lines) between Spike and antibody is Å. **(G)** Substituted residues of Spike involved in the binding interface and their relative energy changes across four predicted models (RRMC, RRMF, AFRC, AFRF). Number of “+” or “-” symbols indicates our confidence in the prediction (three or four: high, two: moderate, one: low), * indicates a situation where there are conflicting predictions from two or more methods.

#### Class 3 Antibodies

Class 3 antibodies bind the RBD in either conformation (up/open or down/closed) at an epitope distal to the ACE2-binding site.^11^ Although their binding sites technically do not overlap with that of ACE2, some may be capable of blocking ACE2 interaction via steric or even allosteric interference. Moreover, they confer the potential for additive neutralization due to their impartiality for RBD conformation and binding sites that do not overlap with Class 1 and Class 2 antibodies.^11^ Examples of Class 3 antibodies include C110, C135, REGN10987, and S309.^11^ We explored the omRBD complexes with single and cocktail therapeutic antibodies currently being administered in the clinical setting.

S309 served as a template for design of an Fc-modified therapeutic antibody, VIR-7831 (sotrovimab; GSK4182136; Xevudy), from Vir Biotechnology and GlaxoSmithKline currently under United States Food & Drug Administration (FDA) Emergency Use Authorization (EUA). S309 was originally isolated from an individual infected with SARS-CoV in 2003 and discovered to also neutralize SARS-CoV-2.^34^ In our structural analysis, G339D and N440K were indicated to be energetically-favorable substitutions in two out of four omRBD-S309 models each (**Fig. 7**). G339D may permit the formation of a novel sidechain interaction while N440K may allow an additional hydrogen bond with its backbone (**Fig. 7A,7B**). As these interactions involve portions of the RBD with lower pTM in AF2 models (**Fig. 2**), greater caution should be taken in drawing direct conclusions. Previous studies of S309, VIR-7831, or VIR-7832 have indicated that G339D and N440K have negligible effects on antibody-binding reduction, ranging from 0.5-1.2-fold reduction.^23,35^ These findings indicate that binding of S309 may be largely unaltered for the Omicron VOC.

**Figure 7:**
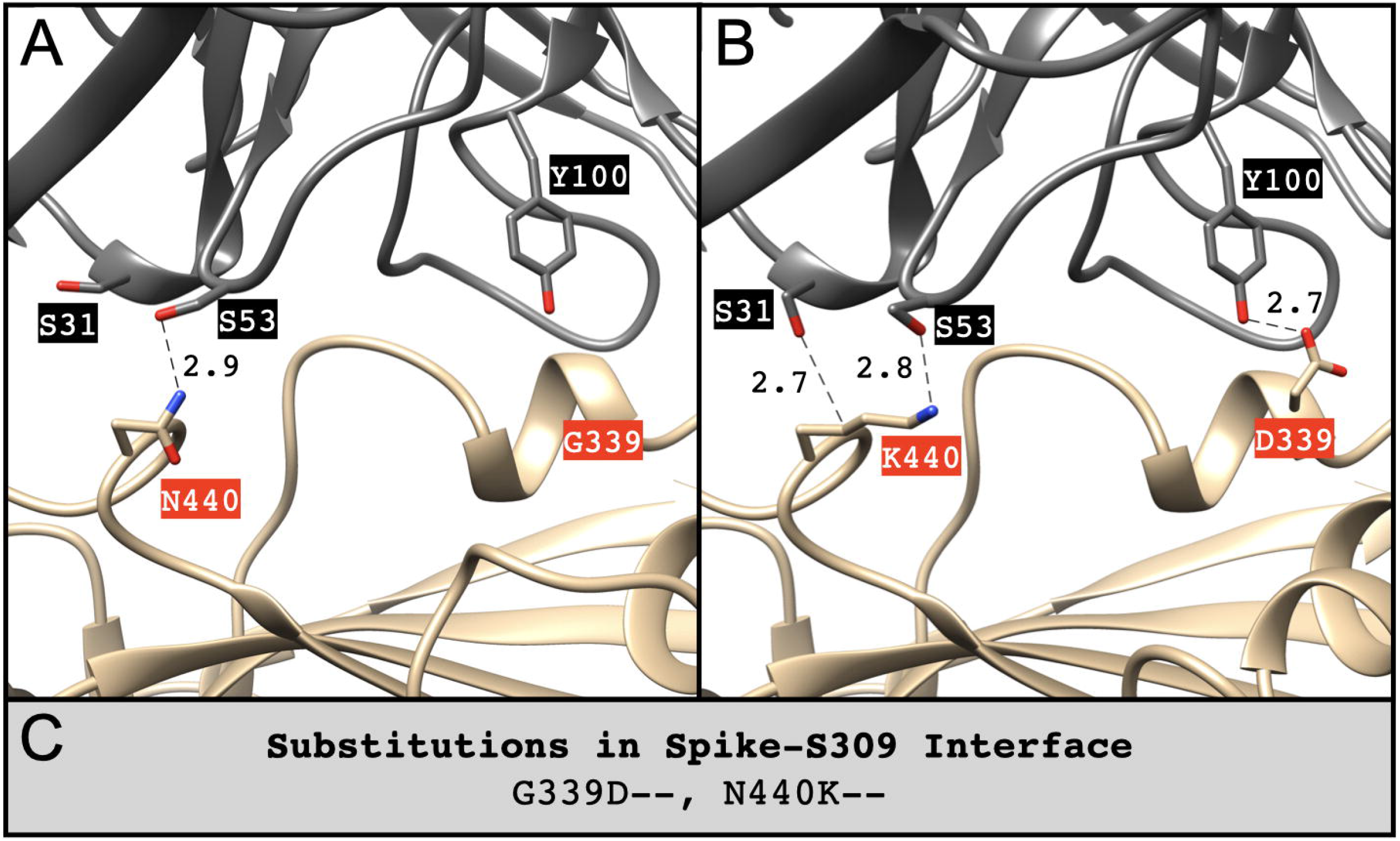
Key substituted residues (red) in the interface between Spike (tan) and S309 (dim gray) that undergo notable energy changes in the Omicron VOC based on its **(A, B)** AFRC model. Length unit of non-covalent bonds (dotted lines) between Spike and antibody is Å. **(C)** Substituted residues of Spike involved in the binding interface and their relative energy changes across four predicted models (RRMC, RRMF, AFRC, AFRF). Number of “+” or “-” symbols indicates our confidence in the prediction (three or four: high, two: moderate, one: low), * indicates a situation where there are conflicting predictions from two or more methods.

REGEN-COV (formerly known as REGN-COV2) from Regeneron Pharmaceuticals is a cocktail composed of two potent antibodies, REGN10987 (imdevimab) and REGN10933 (casirivimab), which are Class 3 and Class 1 antibodies, respectively.^36^ Use of this particular combination highlights the structural advantage of employing antibodies with distinct target epitopes. The cocktail was granted FDA approval for EUA for REGEN-COV in late 2020 and has since provided encouraging phase III clinical trial results.^37,38^ We explored various models of the omRBD-REGN10987 (**Fig. 8A-8E**) and omRBD-REGN10933 (**Fig. 8F-8J**) complexes.

**Figure 8:**
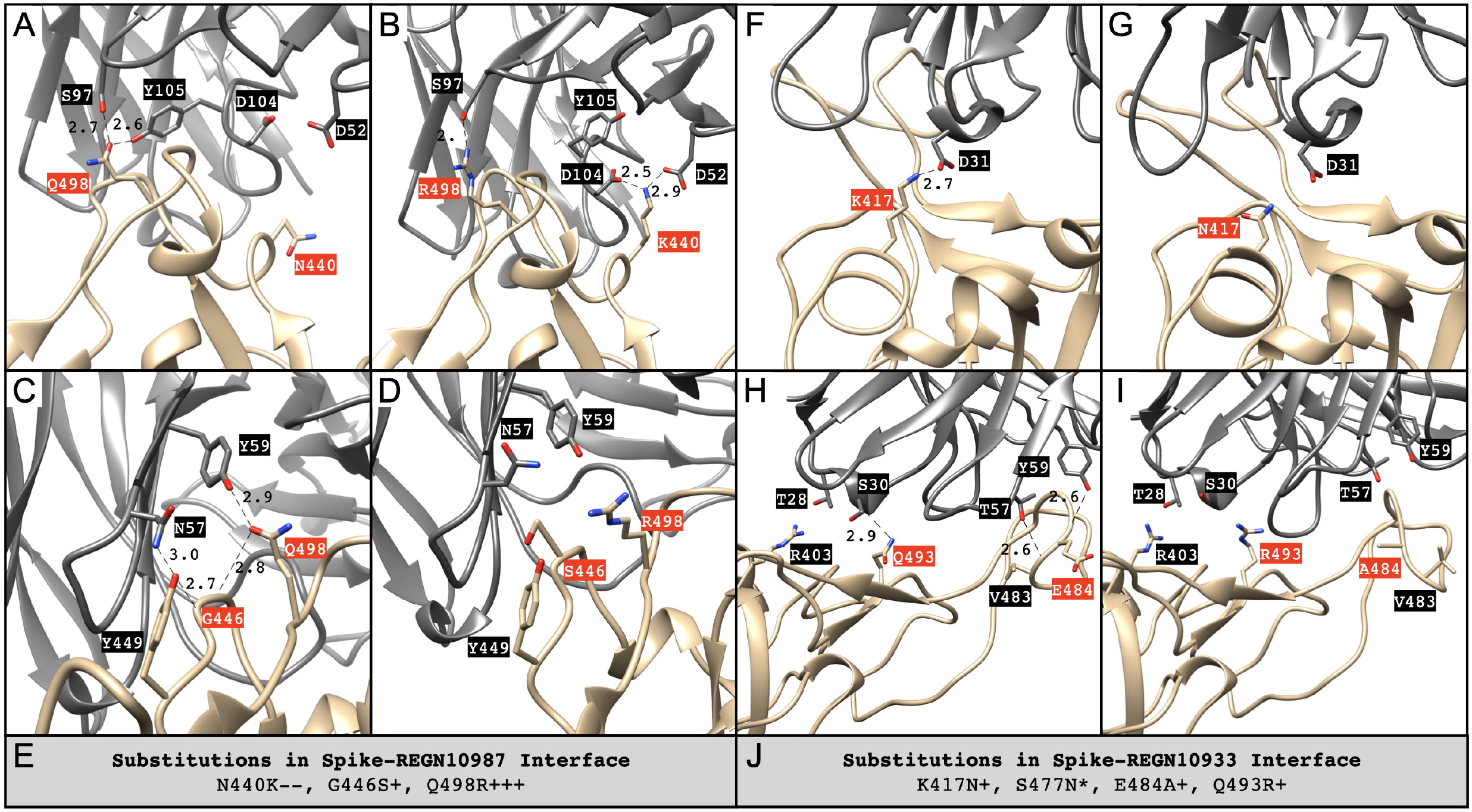
The REGEN-COV cocktail is composed of REGN10987 and REGN10933. Key substituted residues (red) in the interface between Spike (tan) and REGN10987 (dim gray) that undergo notable energy changes in the Omicron VOC based on its **(A, B)** AFRF and **(C, D)** AFRC models (PDB ID: 6XDG). Length unit of non-covalent bonds (dotted lines) between Spike and antibody is Å. **(E)** Substituted residues of Spike involved in the binding interface with REGN10987 and their relative energy changes across four predicted models (RCRC, RCRF, AFRC, AFRF). The same analysis was performed for the interface between Spike (tan) and REGN10933 (dim gray) for its **(F, G)** AFRF and **(H, I)** RCRC models (PDB ID: 6XDG). **(J)** Substituted residues of Spike involved in the binding interface with REGN10933 and their relative energy changes across four predicted models (RCRC, RCRF, AFRC, AFRF). Number of “+” or “-” symbols indicates our confidence in the prediction (three or four: high, two: moderate, one: low), * indicates a situation where there are conflicting predictions from two or more methods.

For omRBD-REGN10987, the data are associated with differing degrees of consensus on energy changes for various residues. Three models reveal a destabilizing impact of Q498R, possibly due to loss of interactions with an adjacent tyrosine residue (**Fig. 8F,8G**). Two models display energetic favorability of N440K, likely due to the formation of two salt bridges with REGN10987 (**Fig. 8F,8G**). Previous studies range greatly in terms of quantified abrogation of binding of REGN10987 in the presence of N440K, ranging from 1.5-to 95.6-fold reduction.^23,39^ Starr et al. found an escape fraction of 0.551 for REGN10987 with N440K.^40^ One model exhibits a positive (*i.e*., unfavorable) energy change associated with S446, which leads to ablation of two backbone hydrogen bonds (**Fig. 8H,8I**) with Y449 and R498. This result is consistent with findings of an escape fraction of 0.795 for G446S by Starr et al.,^40^ and a 17-fold reduction in antibody neutralization of the (chemically similar) substitution Q498H noted in CoVDB.^23^ Overall, our models indicate that G446S and Q498R in omRBD may lead to a loss of interactions (also by Y449) which is in agreement with previous findings with these individual substitutions. However, the (low consensus) prediction of a stabilizing effect of N440K appears to be in contradiction with some previous experimental results.

With regard to REGN10933, some substituted interface residues display unfavorable energy changes; K417N may result in a salt bridge loss while E484A and Q493R may lead to loss of hydrogen bonds. K417N has been shown experimentally to lead to reduction in REGN10933 binding by various degrees, ranging from 4.4-to >100-fold reductions,^29,39,41-45^ while reduced binding by 0.8- to 6.7-fold has been reported for the cocktail with REGN10987.^23,29,39,42^ S477N was reported to lead to 0.9- to 3.4-fold reduction in REGN10933 binding^43,45-47^ and 1.5-fold reduction for the cocktail.^23,46^ Finally, Q493R led to a 70-fold reduction in REGN10933 binding.^23^ In our models, destabilizing energy changes corresponding to these substitutions are observed (**Fig. 8J**), and no compensatory interactions are evident in any of the models, suggesting that the binding to this TE by omRBD may be severely compromised.

#### Class 4 Antibodies

Class 4 antibodies are capable of binding the RBD only in its up/ open conformation at an epitope that does not overlap with the ACE2-binding site.^11^ Examples include CR3022, COV1-16, EY6A, S304, and S2A4.^11^ We analyzed the omRBD-CR3022 complex as a primary example of Class 4 antibody interaction with RBD.

CR3022 was first isolated in 2006 from convalescent sera of a patient infected with SARS-CoV.^48^ It primarily binds to the Wuhan-Hu-1 RBD via a series of hydrophobic interactions.^49^ Of all the identified SARS-CoV-2 Spike-binding proteins in the PDB, CR3022 occurred in the largest number of PDB IDs (**Table 1**). Overall, S373P and S375F appear to contribute favorable and unfavorable energetics, respectively (**Fig. 9**); S373P promotes adoption of an α-helical conformation (which likely influences the structural uncertainty for this region) while S375F eliminates its hydrogen bond with the polypeptide chain backbone (**Fig. 9A,9B**). Such findings indicate that the amino acid substitutions found in the Omicron VOC may produce a neutral outcome with respect to CR3022 binding. Because this binding mode involves lower pTM in AF2 models (**Fig. 2**), we anticipate further testing our methodology in future work by taking advantage of the availability of 14 other PDB structures that involve this antibody, providing some conformational diversity beyond that which we have generated with the approach reported herein.

**Figure 9:**
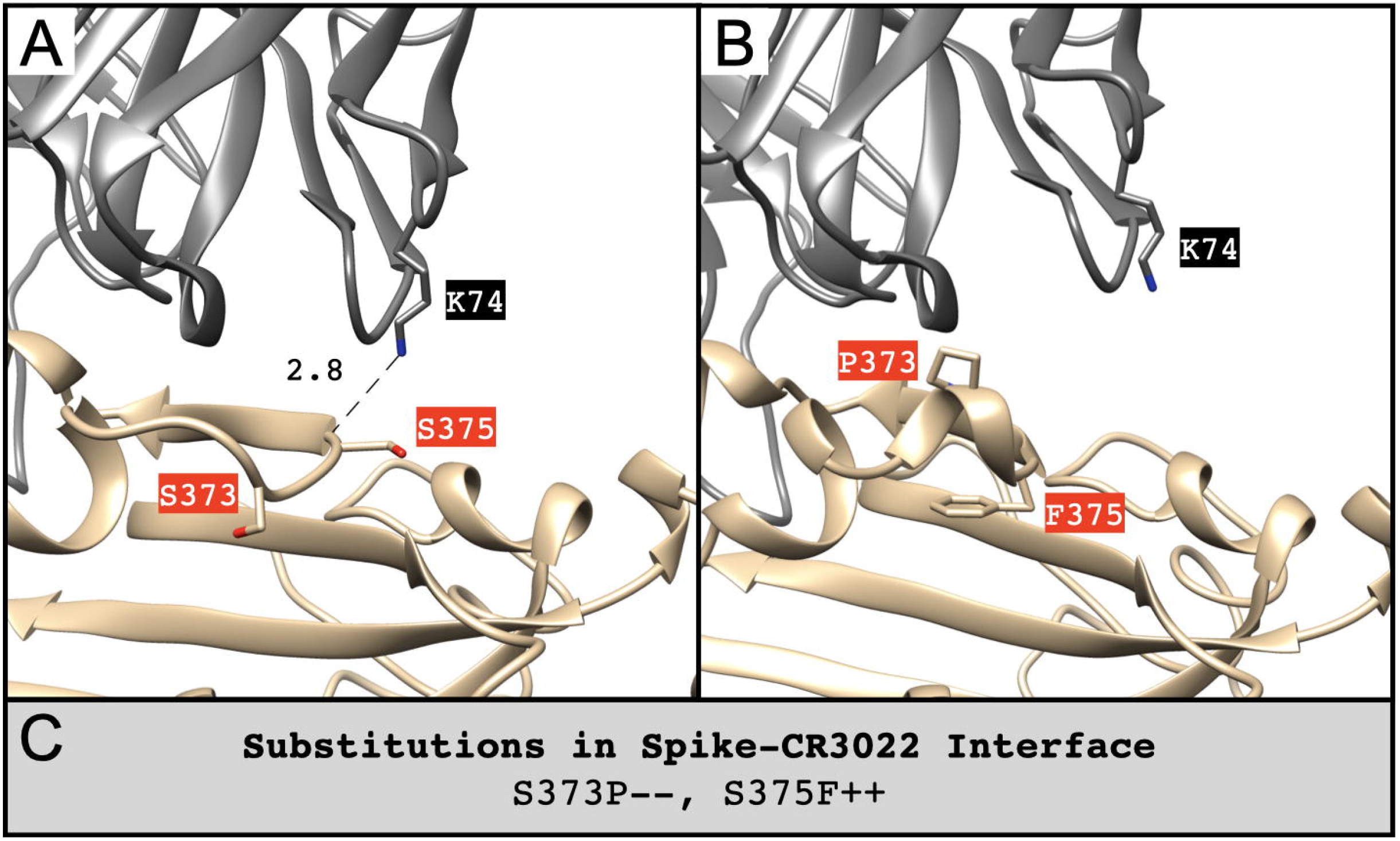
Key substituted residues (red) in the interface between Spike (tan) and CR3022 (dim gray) that undergo notable energy changes in the Omicron VOC based on its **(A, B)** AFRC model (PDB ID: 6YLA). Length unit of non-covalent bonds (dotted lines) between Spike and antibody is Å. **(C)** Substituted residues of Spike involved in the binding interface and their relative energy changes across four predicted models (RRMC, RRMF, AFRC, AFRF). Number of “+” or “-” symbols indicates our confidence in the prediction (three or four: high, two: moderate, one: low), * indicates a situation where there are conflicting predictions from two or more methods.

## Discussion

The emergence and spread of the Omicron VOC of SARS-CoV-2 has raised concerns regarding transmissibility, severity of disease, and susceptibility to previously generated antibodies. To bolster research efforts to characterize this new VOC, we computed structural models of omRBD bound to ACE2 and 20 diverse antibodies using computational methods, including AF2-based structure prediction and Rosetta-based energy calculations. Based solely on these models, we assessed the qualitative impact of substituted residues within the interface between omRBD and various binding partners to facilitate greater understanding of how these substitutions may impact Omicron VOC infection and antibody neutralization.

Analysis of omRBD interactions with various proteins sheds light on how the relatively large number of residue substitutions in this globally emerging SARS-CoV-2 variant may exert their effects individually and display cooperativity. ACE2 binding by omRBD appears to involve an interdigitated network of interactions that was not observed in the Wuhan-Hu-1 strain, suggesting a structural basis for cooperativity (or positive epistasis; **Fig. 3**). Binding of omRBD to examples of Class 1 and 2 antibodies that we analyzed appears to be considerably altered with destabilizing interactions dominating over a small number of stabilizing interactions arising from the numerous substitutions in the Omicron VOC. On the other hand, interaction networks in Class 3 antibodies such as S309 and Class 4 antibodies such as CR3022 appear to be only modestly affected. As initial results, we have presented computed structural models and analyses for 20 binding proteins from a possible total of 281. In due course, we and others will be able to update our results with analyses of the full set of complexes exemplified in the PDB. A comprehensive, if qualitative, picture of the impact of omRBD substitutions on the binding properties of SARS-CoV-2 Spike RBD should hopefully emerge from doing so.

We anticipate that the structural models of omRBD complexes that we have made available will be useful for other research efforts worldwide. First, for wet laboratory investigators interested in characterizing the impact of individual or clusters of substitutions observed in omRBD, the models provide visual aids plus testable hypotheses. Networks of interactions identified for ACE2 binding, for example (**Fig. 3**) can be tested by using site-directed mutagenesis. In cases where binding of therapeutic antibodies is functionally compromised, these models may provide starting points for computer-aided antibody design efforts. Thus, the models reported herein provide atomic level insights into ligand-directed 3D recognition of omRBD by various binding proteins and therapeutic entities.

Several limitations and caveats apply to our computed structural models and analyses. First, post-translational modifications such as glycosylation, which may often be a key element of Spike structure, are not considered in our protein-only models. Many experimental structure determinations also exclude these glycans, and multiple X-ray structures show that the ACE2-binding interface does not explicitly involve glycans. However, some glycosylation patterns do affect binding of certain antibodies. Second, we do not include explicit interfacial water molecules in our modeling procedures. There were no major water-mediated interactions observed experimentally in the complexes we analyzed in detail. However, it is conceivable that new water-mediated interactions have been missed. Both limitations can be addressed by using the models reported here as starting points for more computationally-expensive modeling approaches (such as molecular dynamics simulations) with appropriately glycosylated amino acid sidechains.^50^

In addition to these limitations inherent to modeling procedures, there are well-known limitations of the energy functions used for scoring and energy evaluations plus the limited conformational sampling that is performed in the interest of computational efficiency. Because the overall structure of omRBD is likely to be highly similar to the Wuhan-Hu-1 RBD, limited sampling around the conformations observed in experimentally-derived structures represents a reasonable time-saving approximation. We do, however, explore alternative conformations by using both AF2- and experimentally-derived structures to generate small ensembles of similar structures as starting points of our analyses, and assign confidence based on consensus within the ensemble. Thus, we have hereby provided what we consider a reasonable balance between conformational sampling and computational expediency, though we fully acknowledge that calculated binding energies are highly sensitive to conformation.

The enthalpy of the bound state is but one of the relevant thermodynamic quantities involved in binding. Furthermore, we have not exhaustively sampled the structural ensembles. Given these limitations, we believe it is crucial to not draw premature conclusions about complex biological phenomena such as escape from antibody neutralization, let alone formulate incautious predictions about immune response to the Omicron VOC based on similar analyses (albeit of a smaller number of complexes). Finally, other than ACE2, we do not explore additional identified receptors of Spike, such as TMPRSS2.^51,52^ The structures and predictions provided in this work should serve as a supplement and guide for other researchers studying the mutational consequences of Omicron and other as yet uncharacterized SARS-CoV-2 VOCs.

In summary, we have organized relevant PDB data and generated computational models of omRBD complexes with ACE2 and a diverse set of antibodies from all four Barnes Classes. Based on these models, we observe strikingly altered interface topologies in the computed protein-protein interfaces. We made qualitative inferences about the energetic impact of substituted residues at the interfaces to provide hypotheses regarding their effects on abrogation and structural remodeling of interactions. We make these data available (https://github.com/sagark101/omicron_models) for the broad scientific community to complement ongoing research of the Omicron VOC, and hope that they will be of use to experimental hypothesis testing, therapeutic antibody design, and other efforts to aid the global community’s response to the SARS-CoV-2 pandemic. Future studies will center on generating models based on additional complex structures available in the PDB, and further characterization of their structural features.

## Acknowledgements

This work was supported by Longenbaugh-Levy Donor-Advised Fund research awards (to R.P. and W.A.), National Science Foundation (NSF) grant CBET-1929237 (to S.D.K.), and National Institutes of Health (NIH) grant GM132565 (to S.D.K.). RCSB PDB is jointly funded by the NSF (DBI-1832184), the US Department of Energy (DE-SC0019749), and the National Cancer Institute, National Institute of Allergy and Infectious Diseases, and National Institute of General Medical Sciences of the NIH under grant R01GM133198.

## Author Contributions

R.P., W.A., S.K.B., and S.D.K. conceptualized and conceived the study. J.H.L. and C.M. retrieved and organized relevant PDB data. J.H.L. and D.B. developed computational approaches for generating structural models. J.H.L., C.M., D.B., and S.D.K. analyzed structural models. C.M. visualized structural models. J.H.L., C.M., D.B., and S.D.K. wrote the manuscript, with subsequent editorial contributions from all other authors.

