## Supplementary Materials for "Structural models of SARS-CoV-2 Omicron variant in complex with ACE2 receptor or antibodies suggest altered binding interfaces"

**Model Trimming**. PDB models for antibodies in SAbDab[1] were trimmed to a single Fab (residues up to 120) and the bound region of the SARS-CoV-2 Spike protein (hereafter Spike). The Spike regions were defined as N-terminal domain (NTD): residues 14-305; receptor-binding domain (RBD): residues 319-541. Determination of whether to include one or both regions in the trimmed model were based on the detection of interfacial residues contacted by the complementarity-determining region (CDR) loops of the antibody, as defined in SAbDab. Detection of interfacial residues defined a Spike residue as interfacing with the CDR if the Cα atom is within 5.5 Å of any CDR $C_\alpha$, or within 9 Å of any CDR $C_\alpha$ and the $C_\alpha$-$C_\beta$ vector of that residue pair is within 75°. Trimmed models could include two Spike chains if that contact was specified in SAbDab. For uniformity, all chains were renamed so that Spike chains were A (and B, if multiple Spike chains were present) and antibody chains were H and L as identified in SAbDab. Original chain names are listed in supplementary table pdb_complexes.csv. The trimmed models were considered sufficient to model the energetic consequences at the binding interface, while removing significant amounts of computational optimization that would not elucidate those consequences. All trimmed models were minimized using the Rosetta FastRelax protocol[2] with coordinate constraints restricting backbone movement prior to other modeling steps explained below. Ten decoys were generated and the single lowest-scoring one was used.

**Rosetta Repack/Minimization Modeling**. Repack/minimization models were produced using a FastRelax protocol. The backbone was mobile, and the sidechains of all residues that are mutated in Omicron Variant of Concern (VOC), as well as all interfacial residues (detected as described in the previous paragraph) with them, were optimized. In the wild-type (WT) (*i.e.*, Wuhan-Hu-1) model, the sidechains were optimized with their original sequence in the PDB. In the case of the Omicron VOC (OM) models, the sidechains that were optimized were those corresponding to the mutated sequence. This modeling was done in two ways. In the case of constrained models, a score penalty was applied to inhibit significant Cα movement from the starting model. In the free models, no such constraints were applied, and the optimization scoring was determined purely based on the Rosetta default energy function.[3] Ten decoys were generated and the single lowest-scoring one was used in analyses.

**AlphaFold2 Modeling**. To produce AlphaFold2 (AF2) Spike-antibody structures, the best-ranked WT and OM RBD (omRBD) models were superimposed with the RBD in relaxed trimmed models, and PDBs were generated using the existing antibody structure with the exchanged RBD. Models were then re-minimized using FastRelax, again both with and without constraints. Ten decoys were generated and the single lowest-scoring one was used in analyses.

**Energy Calculations**. Eight models for each Spike-antibody complex, between the two modeling methods, Rosetta repack+minimize (RRM) and AlphaFold2+Rosetta FastRelax (AFR), each used both with and without positional restraints, modeling both WT and OM sequence Spikes. Interfacial energies were calculated as the sum of pairwise energies across an interface (either single-residue with antibody or full Spike structure with antibody).

**Supplementary Table 1**: Benchmark comparisons between the AF2-predicted full-sequence omRBD and several other wild-type Spike structures in complexes.

| PDB ID | $C_\alpha$ RMSD (Å) |
|---|---|
| 6VXX | 0.56 |
| 6XC4 | 0.54 |
| 6XCM | 0.81 |
| 6XDG | 0.90 |
| 6YLA | 0.43 |
| 6ZDH | 0.57 |
| 6ZGE | 0.81 |
| 7BEP | 0.41 |
| 7K8S | 0.66 |
| 7K8X | 0.71 |
| 7KMG | 0.37 |
| 7LRT | 1.01 |
| 7M7W | 0.50 |
| 7MM0 | 1.29 |
| 7NX6 | 0.54 |
| 7ORA | 0.34 |
| 7R6X | 0.40 |

**Supplementary Table 2**: Benchmark comparisons between the AF2-predicted full-sequence Omicron Spike monomer and several variant Spike structures in the PDB.

| PDB ID | $C_\alpha$ RMSD (Å) |
| --- | --- |
| 7EKF (Alpha) | 0.38 |
| 7EKF (Beta) | 0.36 |
| 7EKC (Gamma) | 0.37 |
| 7V89 (Delta) | 0.79 |
| 7NXC (P1) | 0.41 |
| 7BH9 (Evolved RBD) | 0.52 |

**Supplementary Table 3**: Benchmark comparisons between the RBD of the AF2-predicted Omicron VOC Spike monomer RBD and the RBD structures in the antibody-bound complexes used in our analyses.

| PDB ID | Complex | WT $C_\alpha$ RMSD (Å) | OM $C_\alpha$ RMSD (Å) |
|---|---|---|---|
| 6M0J | ACE2 | 1.133 | 1.161 |
| 6CX4 | 1 | 0.601 | 0.611 |
| 6XC4 | 2 | 0.647 | 0.634 |
| 6XCM | 1 | 0.997 | 1.044 |
| 6XCM | 2 | 0.901 | 0.99 |
| 6XDG | 1 | 1.027 | 1.073 |
| 6XDG | 2 | 1.25 | 1.368 |
| 6YLA | 1 | 0.724 | 0.709 |
| 6YLA | 2 | 0.701 | 0.688 |
| 6ZDH | 1 | 0.943 | 1.033 |
| 6ZDH | 2 | 0.912 | 0.903 |
| 6ZDH | 3 | 0.74 | 0.712 |
| 7BEP | 1 | 0.607 | 0.652 |
| 7BEP | 2 | 0.667 | 0.815 |
| 7BEP | 3 | 0.606 | 0.7 |
| 7BEP | 4 | 0.636 | 0.719 |
| 7K8S | 1 | 1.246 | 1.256 |
| 7K8S | 2 | 1.448 | 1.47 |
| 7K8S | 3 | 1.233 | 1.199 |
| 7K8X | 1 | 1.814 | 1.659 |
| 7K8X | 2 | 0.928 | 0.89 |
| 7KMG | 1 | 0.596 | 0.73 |
| 7KMG | 2 | 0.659 | 0.783 |
| 7LRT | 1 | 1.462 | 1.4 |
| 7M7W | 1 | 0.757 | 0.867 |
| 7M7W | 2 | 0.728 | 0.825 |
| 7M7W | 3 | 0.653 | 0.697 |
| 7M7W | 4 | 0.685 | 0.715 |
| 7MM0 | 1 | 2.159 | 2.18 |
| 7NX6 | 1 | 0.611 | 0.653 |
| 7NX6 | 2 | 0.618 | 0.644 |

| | | | |
|---|---|---|---|
| 7ORA | 1 | 0.514 | 0.516 |
| 7ORA | 2 | 0.515 | 0.534 |
| 7ORA | 3 | 0.575 | 0.648 |
| 7ORA | 4 | 0.59 | 0.652 |
| 7R6X | 1 | 0.643 | 0.75 |
| 7R6X | 2 | 0.632 | 0.67 |
| 7R6X | 3 | 0.621 | 0.652 |

**Supplementary Table 4**: antibody_pdb_summary.csv (https://github.com/sagark101/omicron_models/blob/main/antibody_pdb_summary.csv)

PDB entries with identified TEs. PDBs including a cocktail with multiple different antibodies/spike-binding protein types are split into separate rows for each one. The Model Name column identifies the corresponding structures in our trimmed_pdb_structures folder (https://github.com/sagark101/omicron_models/tree/main/trimmed_pdb_structures).

**Supplementary Table 5**: pdb_complexes.csv (https://github.com/sagark101/omicron_models/blob/main/pdb_complexes.csv)
Analysis of the individual antibodies (only antibodies, not nanobodies/sybodies or other, since SAbDab sequences were necessary for CDR and chain identification) in Supplementary Table 4 used for creating the trimmed models. antibody_number identifies what model number each antibody was assigned. Antigen chains were all renamed to A (and B in the case of multi-antigen models) in the trimmed structure, and heavy and light chains were changed to H and L. spike_residues lists the antigen residues interfacing with the antibody CDR loops. in_ntd, in_rbd, in_ace2bind all indicate whether any of the contacted Spike residues were in those regions, with NTD being defined as residues 14-305, RBD as 319-541, and ACE2 binding as 438-506. Multichain indicates whether there are two Spike chains included in the trimmed model. missing_antigen_res lists areas of non-occupancy in the crystal structures. In some cases, mutations occurred in such regions, meaning that those mutations (and their energetic consequences) would not be modeled. We therefore attempted to select representative antibodies without significant density gaps.

**Supplementary Table 6**: energy_and_changed_residue_identification.csv (https://github.com/sagark101/omicron_models/blob/main/energy_and_changed_residue_identification.csv)

Energetic analysis for the binding interfaces. The first four columns identify the antibody and modeling method. wt_total, om_total, and d_total represent the computed Rosetta total energy for the full complex of the Wild Type and Omicron models, and the difference between them. wt_interface, om_interface, and d_interface represent the same, but just the sum of pairwise interfacial scores, rather than the full complex scores. d_interface was used for consensus scoring. Subsequent columns list identified residues with d_total and/or d_interface scores with an absolute value greater than 1 REU. tot_= total energy; inter_ = interfacial energy. stab_ = stabilizing, i.e. d_[energy] < -1 REU; destab_ = destabilizing, i.e. d_[energy] > +1 REU. mut_ = mutated residues; other_ = sites conserved in Omicron. close = in the region of the Spike structure that interfaces with the antibody; distal = not in the region of the Spike structure that interfaces with the antibody. Interface interactions were only in the close regions, as distal residues did not influence interface score.

**Supplementary Table 7**: single_res_energies.csv (https://github.com/sagark101/omicron_models/blob/main/single_res_energies.csv)

Computed energies for each residue in each analyzed antibody-Spike complex. The first four columns identify the antibody and modeling method. The _tot and _inter columns are the same as those described in Table 6, though on a single-residue basis rather than for the full structure, i.e the sum of pairwise interactions a given Spike residue has with all antibody residues. The _intra energy columns are the computed intra-chain energies of the residues, i.e. the interfacial energy a given Spike residue has with all Spike residues other than itself.

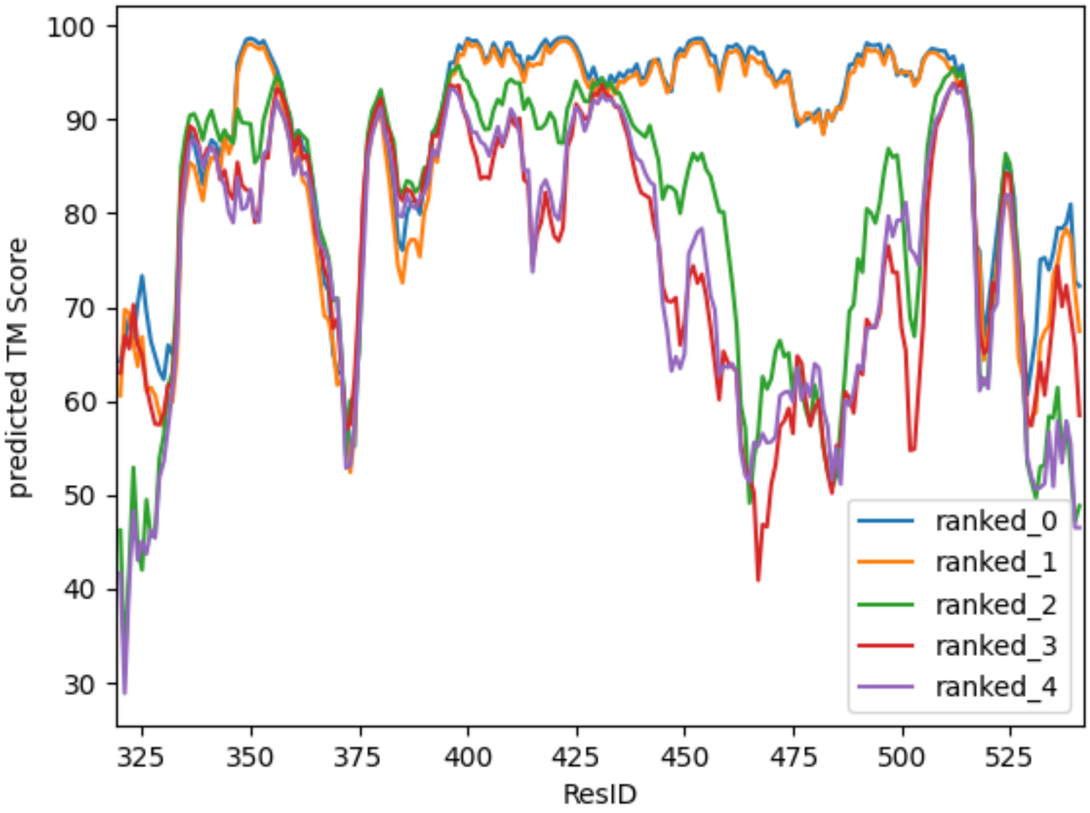


**Figure S1**: pTM of each residue within omRBD for the five AF2-predicted models. Residues (ResIDs) of isolated omRBD are numbered according to their positions relative to wild-type Spike.
